# Resistance to Carcinogenesis in the African Spiny Mouse (*Acomys*) correlates with upregulation of tumor suppressor genes

**DOI:** 10.1101/2024.10.24.620065

**Authors:** Marta Vitorino, Gonçalo G. Pinheiro, Ines M. Araujo, Bibiana I. Ferreira, Wolfgang Link, Gustavo Tiscornia

## Abstract

Cancer remains a leading cause of morbidity and mortality worldwide, driving extensive research into the mechanisms that contribute to its development and progression. The search for animal models capable of shedding light on mechanisms of cancer resistance has led researchers to explore various rodent species. Among these, the African Spiny Mouse (*Acomys sp.)* has attracted considerable attention due to its regenerative abilities. In our study, we compared the response of *Mus musculus* and *Acomys dimidiatus* mice to treatment with the carcinogen 7,12-dimethylbenz[a]anthracene (DMBA) combined with the inducer of proliferation 12-O-tetradecanoylphorbol-13-acetate (TPA). While both *Acomys* and *Mus* mice experienced carcinogenic damage to their skin cells, mounted proliferative responses and underwent immune cell infiltration, only *Mus* mice developed tumors, whereas *Acomys* remained tumor-free. To uncover the molecular mechanisms underlying this resistance, we performed RNA sequencing on tissue samples from both species at baseline and at multiple time points during the carcinogenesis protocol. The data reveal distinctly different transcriptional responses between both species. In particular, *Acomys* showed massive upregulation of immune related genes, including a set of tumor suppressor genes, while *Mus* seemed to focus its response on modifying epidermis structure and regulation of the cell cycle. These differences correlated with an increased apoptotic response in *Acomys* but not *Mus*. In summary, our study provides compelling evidence that *Acomys* mice exhibit remarkable resistance to tumor development under DMBA/TPA carcinogen treatment mediated through the upregulation of a range of processes related to immune surveillance.

## Introduction

The relationship between cancer, wound healing and regeneration is complex and unresolved. Upon injury, after an initial response to achieve hemostasis, organisms have two fundamental choices: regeneration or wound healing. Regeneration involves recreation of the original tissue architecture, while wound healing typically involves fibrotic scarring (Krafts, 2010). Regeneration involves a mandatory proliferative response designed to provide the cellular mass necessary for tissue reconstruction, which, if not strictly regulated, can lead to uncontrolled proliferation and cancer. A critical phase of wound healing is re-epithelization, where keratinocytes adjacent to the wound hyper- proliferate and migrate into the wound bed, both behaviors very similar to what occurs in cancer initiation and metastasis. In regeneration, proliferation is orderly, and importantly, responds to termination signals; if these signals are overridden, cancer may ensue. Furthermore, regeneration, wound healing and cancer share major underlying molecular mechanism and pathways (Maggiore & Zhu, 2024; Rybinski, 2014; Schäfer & Werner, 2008) Arnold et al., 2015; Foster et al., 2018; Sundaram et al., 2018; Ganesh et al., 2020),to the point where it has been proposed that cancer is a wound that never heals (Dvorak, 2015; Flier et al., 1986). The relationship between regeneration and cancer is reinforced by observations throughout the animal kingdom. One prevailing hypothesis suggests that high regenerative capacity tends to be associated with a low susceptibility to cancer, presumably due to trade-offs between both outcomes(Wong et al., 2020). Planarians are capable of regeneration of entire organs even from small body fragments (Reddien, 2018), a process driven by pluripotent stem cells known as neoblasts (Rink, 2013); however, the extent of this regenerative capacity varies across planarian species. Planarians from the genus *Schmidtea* are capable of full body regeneration, while those in the genus *Dugesia* exhibit more restricted regenerative abilities. Interestingly, *Dugesia* species also show a tendency to develop spontaneous tumors (Stephan, 1962)as well as after exposure to cadmium (Hall et al., 1986; Voura et al., 2017), while *Schimidtea* does not, suggesting a link between reduced regenerative capacity and increased cancer susceptibility (Plusquin et al., 2012). In *Schmidtea*, knock-down of the planarian ortholog of p53 results both in tumorigenesis and impaired regeneration (Oviedo et al., 2008; Pearson et al., 2010). Urodeles, including axolotls, salamanders, and newts, are capable of regenerating multiple tissues and organ systems. Despite axolotls having been observed to spontaneously develop skin tumors (Harshbarger et al., 1999), urodeles are thought to have low rates of tumorigenesis, and are resistant to cancer after carcinogen exposure (Ingram, 1971). Zebrafish can regenerate a range of tissues after injury (Gemberling et al., 2013). While spontaneous tumor formation in zebrafish is rare (Liu et al., 2011; White et al., 2013) exposure to carcinogens induces vigorous tumor formation in these animals. The power of the genetic toolbox available for this model has led to a number of studies linking cancer susceptibility to regenerative capability (Demirci et al., 2022; Hale et al., 2017).

Intriguingly, cancer incidence varies widely across the animal kingdom, with certain species in every taxon exhibiting remarkably low rates of tumorigenesis(Albuquerque et al., 2018). Possible mechanisms hypothesized to be involved include lower somatic mutation rate, shorter telomeres, redundancy of tumor suppressors, a more efficient immune system, higher apoptosis rate and increased contact inhibition, among others (Caulin et al., 2011). Mechanisms that could explain low cancer incidence rate in non-traditional mammalian models are of great interest. Species with naturally low cancer incidence such as marsupials, bats, elephants, and whales present valuable opportunities for cancer research, but pose significant logistical challenges as research models (Albuquerque et al., 2018). Naked mole rats offer a more convenient model but come with specific housing and care needs due to their eusocial nature (Ragland et al., 2022). Although common laboratory rodents are easy to breed, their high cancer rates limit their usefulness for studying cancer resistance (Greenacre, 2004). Therefore, a rodent model with low cancer incidence would be particularly valuable to investigate natural cancer resistance mechanisms. In this study, we investigated the susceptibility of the African spiny mouse (*Acomys dimidiatus*), an emerging non-traditional rodent model, to chemically induced tumor formation. The most striking feature of *Acomys* is its exceptional regenerative capacity (Seifert et al., 2012; Gawriluk et al., 2016; Matias Santos et al., 2016; Maden et al., 2018; Maden & Brant, 2018; Nogueira-Rodrigues et al., 2022; Okamura et al., 2022;) which led us to establish this species as a research model several years ago. Despite their relatively long lifespan, we have never observed uncontrolled cell proliferation or spontaneous tumor development. This absence of cancer prompted us to systematically explore their cancer resistance. Here we report the resistance of *Acomys* mice to the carcinogen treatment 7,12-dimethylbenz[a]anthracene (DBMA) followed by the proliferation inducer 12-O-tetradecanoylphorbol-13-acetate (TPA). The DBMA/TPA model is a well-established experimental system used to study chemically induced skin carcinogenesis (Li & Brakebusch, 2021; Vähätupa et al., 2019), particularly in *Mus musculus.* Our work shows that *Acomys* dimidiatus is resistant to DMBA/TPA induced tumorigenesis and provides valuable insights into the underlying protective mechanisms operating in *Acomys*.

## Material and Methods

### Animals

Specimens of *A. dimidiatus* were kept at the animal facility of the Algarve Biomedical Center Research Institute at the University of Algarve, in a room with controlled temperature (26°C), on a 13/11 h light/dark cycle and fed twice a week with mixed seeds supplemented with fresh fruits and vegetables, with water ad libitum, as described for this species (Pinheiro et al., 2018). All experiments were performed in accordance with European guidelines (2010/63/EU) for the care and use of laboratory animals, as well as Portuguese law (DL 113/2013). All experimental procedures were reviewed and approved by the Animal Welfare Body of the University of Algarve and have been subjected to approval by the Direcção Geral de Alimentação e Veterinária of Portugal.

### DMBA/TPA treatment

To evaluate the induction of tumors by DMBA/TPA two stage protocol, the dorsal hair on the back of *Mus musculus* C57BL/6N and *Acomys dimidiatus* animals was trimmed and the skin was treated topically with DMBA (Sigma-Aldrich; 100 μg in 100 μL acetone). Starting one week after DMBA application, animals were treated twice weekly with TPA (Sigma-Aldrich; 25 μg in 100 μL absolute ethanol) for 30 weeks. Tumor formation was assessed weekly. For a transcriptomic analysis of events during the first 28 days of treatment, *Mus* and *Acomys* animals were subjected to the same tumor inducing protocol and samples harvested as described above at day 1 (D1, 24 hours after treatment with DMBA), day 14 (D14, after initial treatment with DMBA and TPA treatment on D7, D9 and D11) and day 28 (D28, after initial treatment with DMBA and TPA treatment 3 times a week from D7 to D28).

### Immunohistochemistry

Immunohistochemistry was performed in the Histopathology Core Facility at the Institute for Research in Biomedicine in Barcelona (Spain), following standard protocols. Briefly, *Acomys* dorsal skin was fixed in 4% paraformaldehyde overnight, embedded in paraffin, and cut into 3-5 μm sections. For immunostaining, the sections were dewaxed and epitope retrieval was performed with ER1 buffer (AR9961, Leica Biosystems) for Ki67 (Abcam, 15580) for 30 min and with ER2 buffer (AR9640, Leica Biosystems) for Cleaved-Caspase 3 (Cell Signaling, 9661), CD45 (Cell Signaling, 98819), H2AX (Cell Signaling, 9718) and CD68 (Byorbit, orb47985) for 20 min. Washings were performed using the BOND Wash Solution 10x (AR9590, Leica). Quenching of endogenous peroxidase was performed by 10 min of incubation with Peroxidase-Blocking Solution at RT (S2023, Dako, Agilent). Non-specific unions were blocked using 5 % of goat normal serum (16210064, Life technology) mixed with 2.5 % BSA diluted in wash buffer for 60 min at RT. Secondary antibody used was the BrightVision Poly-HRP-Anti Rabbit IgG Biotin-free, ready to use (DPVR-110HRP, Immunologic). Antigen–antibody complexes were reveled with the DAB (Polymer) (Leica, RE7230-CE). Sections were counterstained with hematoxylin (RE7107, Leica Biosystems) and mounted with Mounting Medium, Toluene-Free (CS705, Dako, Agilent) using a Dako CoverStainer. For Iba1 (019-19741, Wako), samples were dewaxed and antigen retrieval process using citrate buffer pH6 for 20 min at 97°C using a PT Link (Dako – Agilent) was performed. Blocking was performed with Peroxidase-Blocking Solution at RT (S2023, Agilent) and 5 % of goat normal serum (16210064, Life technology) mixed with 2.5 % BSA diluted in wash buffer for 10 and 60 min at RT. The secondary antibody used was the BrightVision poly HRP-Anti-Rabbit IgG, incubated for 45min (DPVR-110HRP, ImmunoLogic). Antigen–antibody complexes were reveled with 3-3′-diaminobenzidine (K346811, Agilent). Sections were counterstained with hematoxylin (CS700, Dako, Agilent) and mounted with Mounting Medium, Toluene-Free (CS705, Agilent) using a Dako CoverStainer. Specificity of staining was confirmed staining with the rabbit IgG, polyclonal (NBP2-24891, Novus bio-tec). Digital scanned brightfield images were acquired with a NanoZoomer-2.0 HT C9600 scanner (Hamamatsu, Photonics, France) equipped with a 20X objective and using NDP.scan2.5 software U10074-03 (Hamamatsu, Photonics, France). All images were visualized with the NDP.view 2 U123888-01 software (Hamamatsu, Photonics, France) with a gamma correction set at 1.8 in the image control panel of the NDP.view 2 U123888-01 software (Hamamatsu, Photonics, France). Cell quantification was performed using QuPath 0.5.0 software, employing the automated Positive Cell Detection tool.

### RNA sequencing

Total RNA was extracted from mouse and *Acomys* skin tissues using Tryzol (Nzytech). Briefly, skin was first homogenized using Navy lysis kit (Next Advance) and extracted using Tryzol (Nzytech). RNA integrity was assessed using the Bioanalyzer 2100 system (Agilent Technologies). Messenger RNA was enriched from total RNA using poly-T oligo-attached magnetic beads. After fragmentation, the first strand cDNA was synthesized using random hexamer primers, followed by the second strand cDNA synthesis using dUTP. Libraries were subjected to end repair, A-tailing, adapter ligation, size selection, amplification, and purification. The libraries were checked with Qubit and real-time PCR for quantification and Bioanalyzer 2100 system (Agilent Technologies) for size distribution detection. After library quality control, different libraries were pooled based on the effective concentration and targeted data amount, then sequenced by Novogene Europe on the NovaSeq™ X Plus platform (Illumina).

Raw data (raw reads) of fastq format were processed through fastp software. In this step, clean data (clean reads) were obtained by removing reads containing adapter, reads containing poly-N and low-quality reads from raw sequence. At the same time, Q20, Q30 and GC content of the clean data were calculated. All the downstream analyses were based on high quality, clean reads. The trimmed reads were mapped to either the *Mus musculus* (GCA_947599735.1) or *Acomys dimidiatus* (GCA_907164435.1) reference genomes using HISAT2 v2.05 software. We obtained over 20,000 reads for each species. Because the *Acomys* genome is incompletely annotated, we limited our downstream analysis to a subset of 16,542 genes which showed an unequivocal 1 to 1 ortholog relationship between both species, using *Mus* related databases to complete the analysis. The mapped reads of each sample were assembled by StringTie (v1.3.3b) (Pertea, 2015) in a reference-based approach. FeatureCounts v1.5.0-p3 was used to count the read numbers mapped to each gene and then FPKM of each gene was calculated based on the length of the gene and reads count mapped to this gene. Prior to differential gene expression analysis, for each sequencing library, read counts were adjusted using the edgeR R package (3.22.5) by scaling normalization factors to eliminate differences in sequencing depth between samples. Differential expression analysis for two conditions/groups was performed using the DESeq2 R package (1.20.0). The resulting p-value was adjusted using the Benjamini and Hochberg’s methods to control the error discovery rate. The corrected p-value ≤ 0.05 & log2 (foldchange)| was set at 1 for D1, and 2 for D14 and D28, as the threshold of significant differential expression. GO terms, Kegg pathways were analyzed using Panther (https://www.pantherdb.org) and NovoMagic (Novogene) bioinformatic software.

## Results

### Cancer resistance in Acomys

Similar to cancer, tissue regeneration starts with a high rate of proliferation to re-construct the damaged tissue. However, unlike cancer, the proliferative process in regeneration is tightly regulated and tumorigenesis is generally not observed. Notably, there are no reports of cancer resistance in *Acomys* in the literature. In over 10 years of research on our *Acomys* colony at the University of Algarve, focusing on skin and ear wound regeneration and involving more than a thousand animals, we have never observed uncontrolled proliferation or the development of spontaneous tumors in aging animals, despite their relative long lifespan of six years. To explore whether *Acomys*’ regenerative capacity involves specific mechanisms of cancer resistance, we employed a well-established two-stage DMBA/TPA epidermal cancer induction protocol (Sandoval et al., 2022; Vähätupa et al., 2019). Animals were treated with an initial dose of DMBA, a mutagen known to cause DNA double strand breaks, followed by regular injections of TPA to induce proliferation, as described in Figure 1A and Materials and Methods. Remarkably, after 30 weeks of the protocol, 5 out of 6 *Mus* (C57/Bl6) developed multiple papillomas in the treated area (Figure 1B and D), whereas 6 out of 6 *Acomys* exhibited no tumor formation (Figure 1C and D). Figure 1E illustrates the average number of papillomas per animal over the course of the DMBA/TPA treatment.

**Figure 1.**
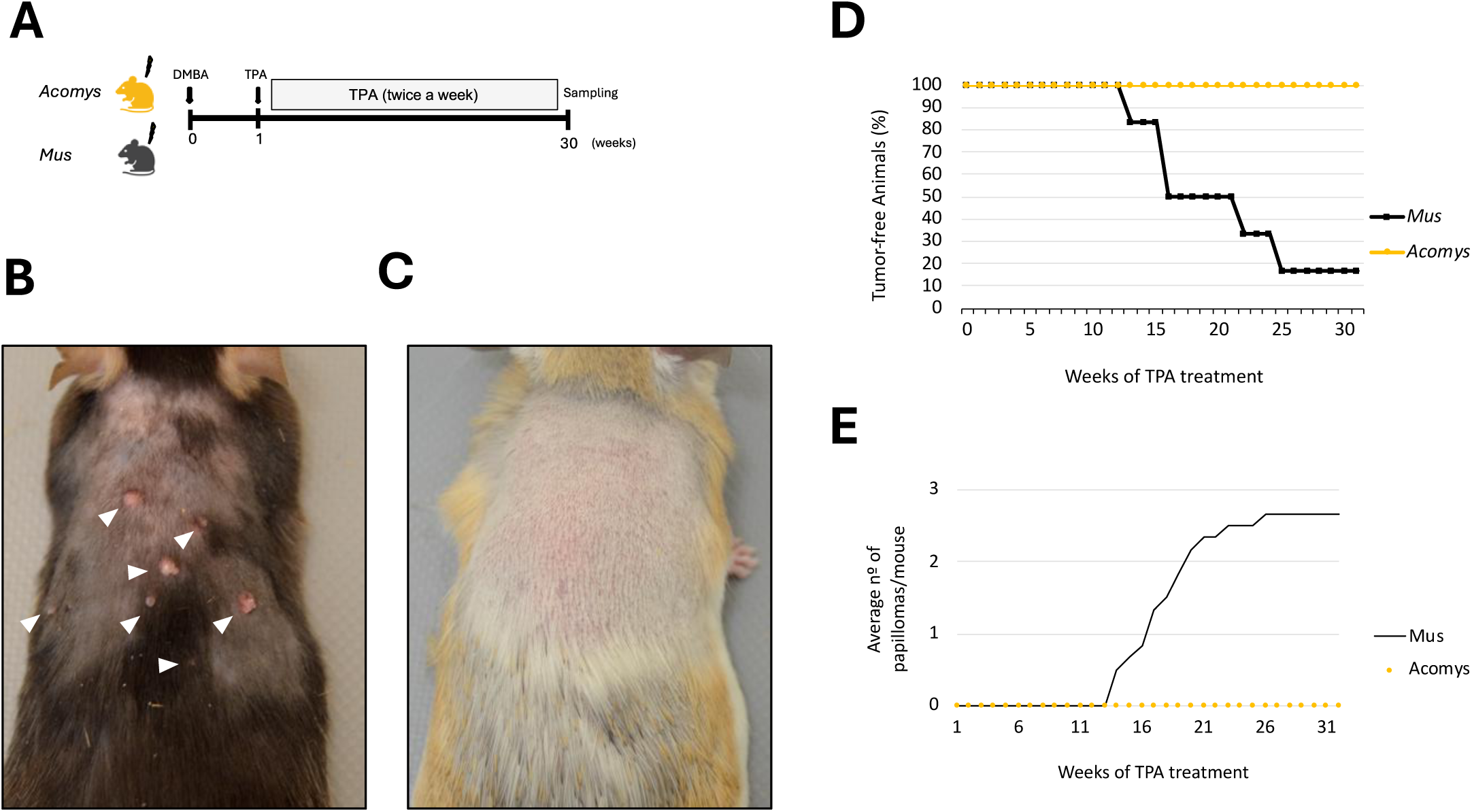
***Acomys dimidiatus* is resistant to tumor development induced by DMBA/TPA two stage protocol**. **(A)** Schematic diagram for carcinogenesis induction by DMBA/TPA treatment on the back skin of both *A. dimidiatus* and *M. musculus*. **(B-C)** Back dorsal skin of *M. musculus* and *A. dimidiatus* 30 weeks after DMBA/TPA treatment with papillomas visible in *M. musculus* **(B)** and absent from *A. dimidiatus* **(C)**. **(D)** Graphical representation of papilloma free animals and (**E**) average number of papillomas per animal over time during DMBA/TPA treatment.

Several mechanisms could explain the strikingly different phenotype observed after 30 weeks of treatment. First, we asked whether the initial DMBA treatment was inducing double strand breaks (DSBs) to similar levels in both species. We found that there was little or no difference in the initial level of genomic insult, as immunohistochemistry (IHC) against the DSB marker p2AHx revealed similar levels of DNA damage in the epidermis of both *Mus* and *Acomys* 24 hours after the initial treatment with DMBA (Figure 2A and B). While DMBA induced DSBs are clearly present within 24 hours of DMBA treatment in both species, it is possible that an enhanced or rapid DNA repair response in *Acomys* resolves the DSBs injury quickly, precluding the establishment of tumorigenic mutations.

**Figure 2.**
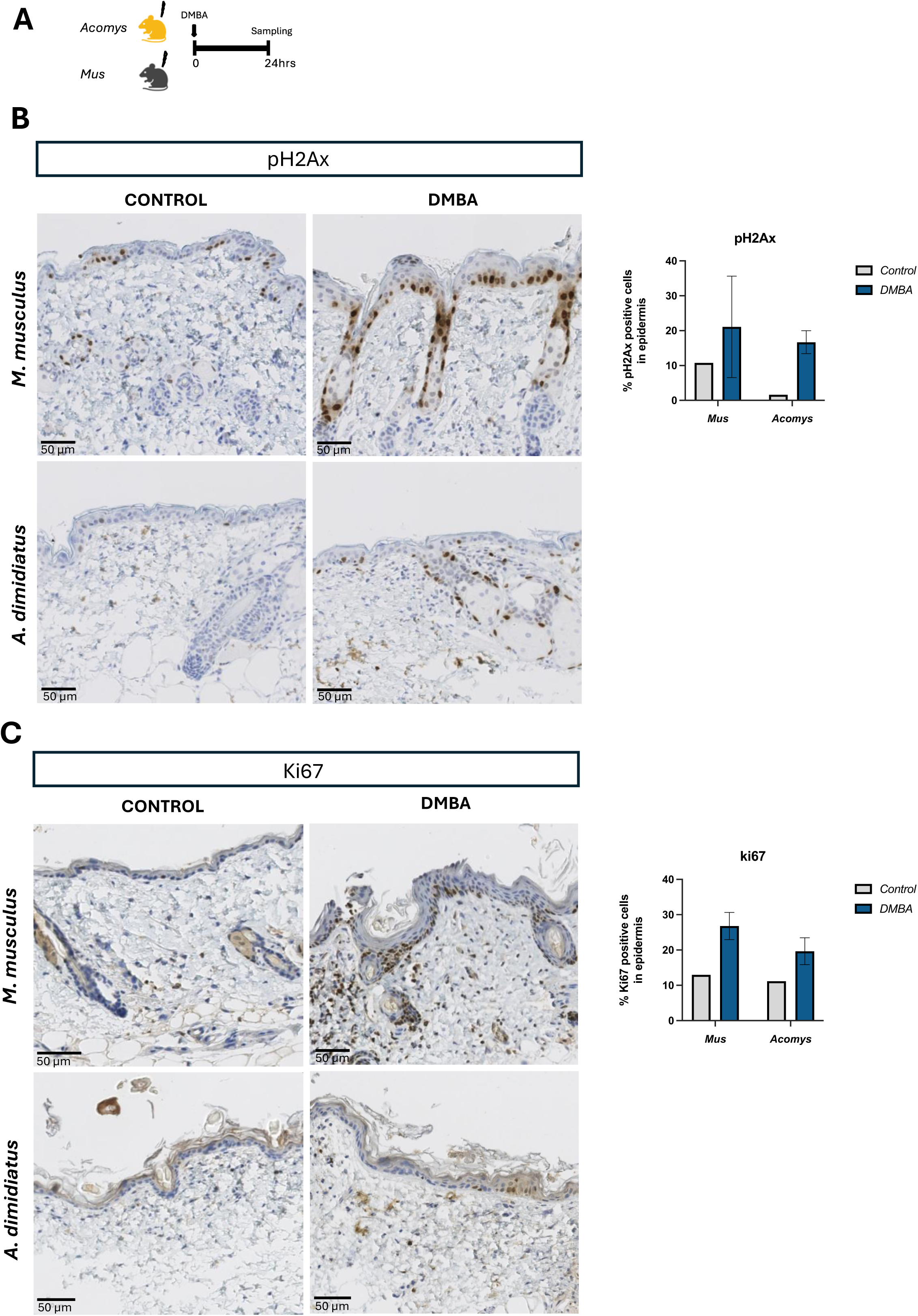
DMBA triggers double-strand-break formation in both *A. dimidiatus* and ***M. musculus.* (A)** Schematic illustration depicting the experimental approach to investigate short-term responses to DMBA treatment of both species. **(B-C)** immunohistochemistry staining and quantification of **(B)** pH2Ax-positive cells and **(C)** Ki67- positive cells in *M. musculus* and *A. dimidiatus* one day after DMBA application. Scale bar: 50 μm.

Concurrently with presence of DSBs, we observed an immediate proliferative response in the epidermis of both species, as indicated by IHC against Ki67, a canonical proliferation marker. Therefore, the initial effect of the DMBA genotoxic insult appear consistent for both species (Figure 2C).

We next aimed to examine whether complete absence of tumors in the African Spiny Mouse could be attributed to early events occurring within the first month after treatment that effectively halt tumorigenesis in *Acomys*. To investigate this, we conducted a transcriptomic analysis at three different time points: 1 day (D1), 14 days (D14) and 28 days (D28) after the application of the carcinogen.

### Transcriptional Profile one day after *DMBA* treatment

Samples from *Mus* and *Acomys* collected at D1 (24 hours after treatment with DMBA) were compared to samples from untreated mice harvested at D0 (N=4, Figure 3A). PCA analysis revealed expected clustering of the samples including the *Acomys* samples, which exhibited some heterogeneity due to the partially outbred nature of the animals of our colony. The PCA1 and PCA2 components explained 41.56% and 18.72% of variation in *Mus* and 41.19% and 22.41% in *Acomys,* respectively (Supplementary Figure 1). We queried differentially expressed genes (DEGs) (log2 =1) and p-adj ≤ 0.05. We comparatively examined genes exclusively upregulated in *Mus* but not in *Acomys* (414) and genes exclusively upregulated in *Acomys* but not in *Mus* (17) (Figure 3B, 3C and 3E). Only 11 genes were upregulated in both species (Figure 3C). To understand what biological processes (BP) were associated with the set of genes upregulated exclusively in each species, we conducted a Panther analysis. Upregulated *Mus* genes resulted in 138 enriched BP ontological categories, the genes of which could be grouped in the following biological themes: cell cycle and proliferation related processes (31.4%); processes related to epidermis structure and morphogenesis (7.1%); processes related to apoptosis (3.2%) and other processes (57.6%). In contrast, no BP ontological categories were significantly enriched for genes exclusively upregulated in *Acomys*, and only 3 BP ontological categories were enriched due to genes upregulated in both species (Figure 3D and 3H).

**Figure 3.**
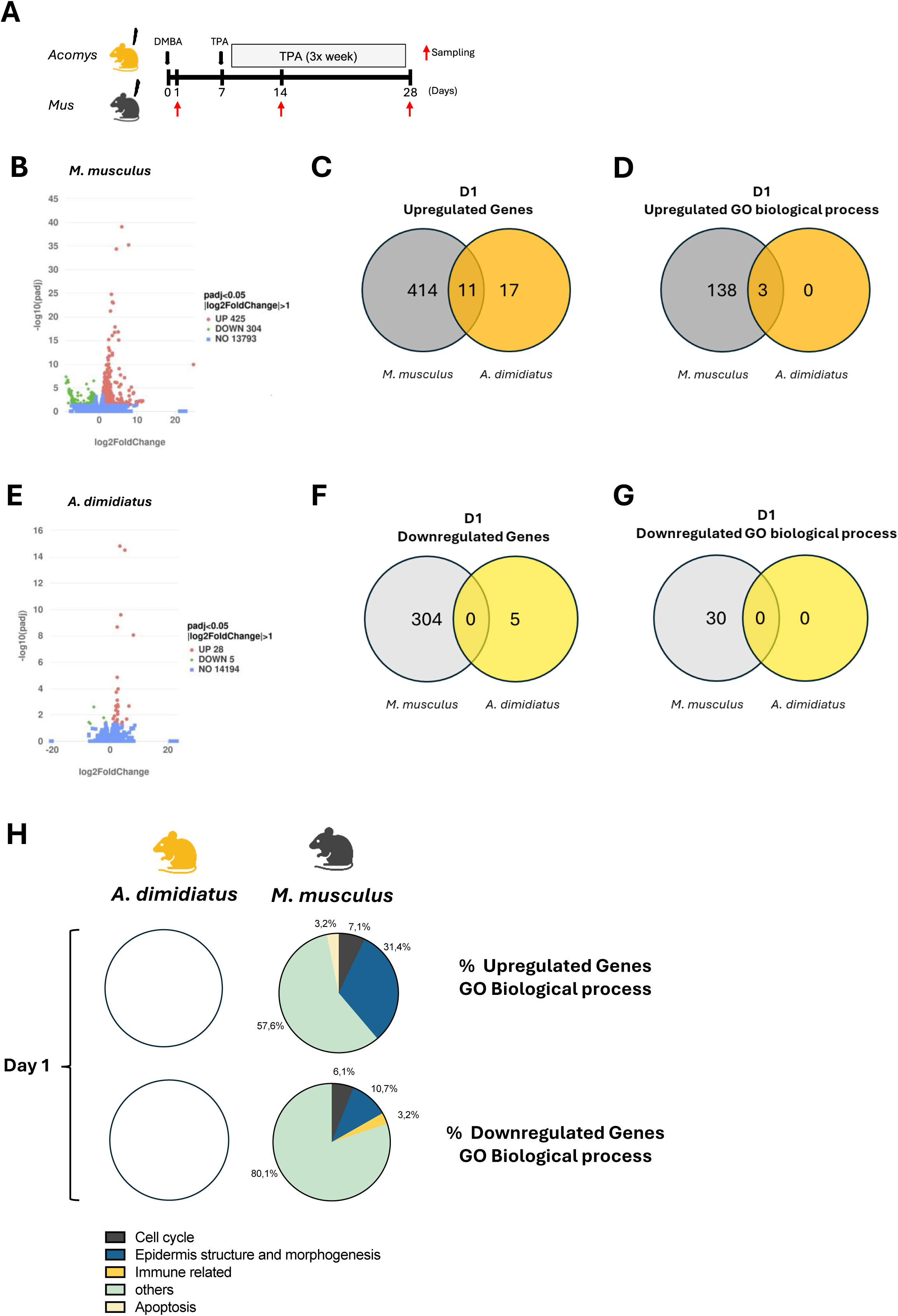
**Administration of DMBA does not induce major differences in the transcriptomic profiles**. **(A)** Schematic illustration of transcriptomic experimental design. Red arrow indicates the times of sample collection. Volcano plots of genes differentially expressed in *M. musculus*. **(B, E)** Volcano plots of genes differentially expressed in *M. musculus* **(B)** and *A. dimidiatus* **(E)** one day after treatment with DMBA (D1). The log2 fold change (FC) indicates the mean expression level for each gene. Genes were scored as differentially expressed when log2 FC>1, p<0,05. Each dot represents one gene. Red dots represent upregulate genes, green dots represent downregulated genes and blue dots represent genes that are not differentially expressed. Venn Diagrams of upregulated genes **(C)** and GO biological processes **(D)**, downregulated genes **(F)** and biological processes **(G)** in both species. **(H)** Percentage of upregulated and downregulated genes included in categories of GO biological processes. White circles are shown when there were no differentially expressed genes identified.

We then examined genes exclusively downregulated in *Mus* but not *Acomys* (304), or exclusively downregulated in *Acomys* but not *Mus* (5) (Figure 3B, 3E and 3F). No downregulated genes were shared between species (Figure 3F). To understand what BPs were associated with the set of genes upregulated exclusively in each species, we performed a Panther analysis. Downregulated *Mus* genes revealed 30 enriched BP ontological categories (Figure 3G), the genes of which could be organized in the following biological themes: cell cycle and proliferation related processes (6.1 %); processes related to epidermis structure and morphogenesis (10.7%); processes related to immunity (3.2%) and other processes (80.1%). In contrast, no BP ontological categories were significantly enriched for genes exclusively downregulated in *Acomys*, or genes downregulated in both species (Figure 3H).

A BP and KEGG pathway analysis conducted with Novomagic (Novogene Europe) showed that response to the DMBA insult at 24 hours did not result in either BP categories or KEGG pathway enrichment involving downregulated genes in either *Mus* or *Acomys.* However, the analysis did identify a number of BP categories and KEGG pathways significantly enriched for upregulated genes in both species (Supplementary Figure 2). Therefore, the initial response to DMBA seems to be mediated by upregulation of specific gene sets in each species. We were particularly interested in the 17 genes upregulated in *Acomys* but not in *Mus*. Among this set we found several genes with intriguing functions related to tumorigenesis. *NQ01*, a NADP dehydrogenase (log2 upregulated 3.45-fold) is involved in detoxification and preventing formation of reactive oxygen species, possibly preventing cellular damage (Oh et al., 2015). Two other genes with tumor suppressor functions exclusively upregulated in *Acomys* are *GNK1* (log2 8.04-fold) and *SPINK7* (log2 4.13-fold). Intriguingly, we found *CYP1A1*, a cytochrome P450 enzyme which is essential for the biotransformation of polycyclic aromatic hydrocarbons such as DMBA (Androutsopoulos et al., 2009) upregulated log2 5.1-fold in *Acomys*, but only log2 3.1-fold in *Mus*) (Table 1).

**Table 1:**
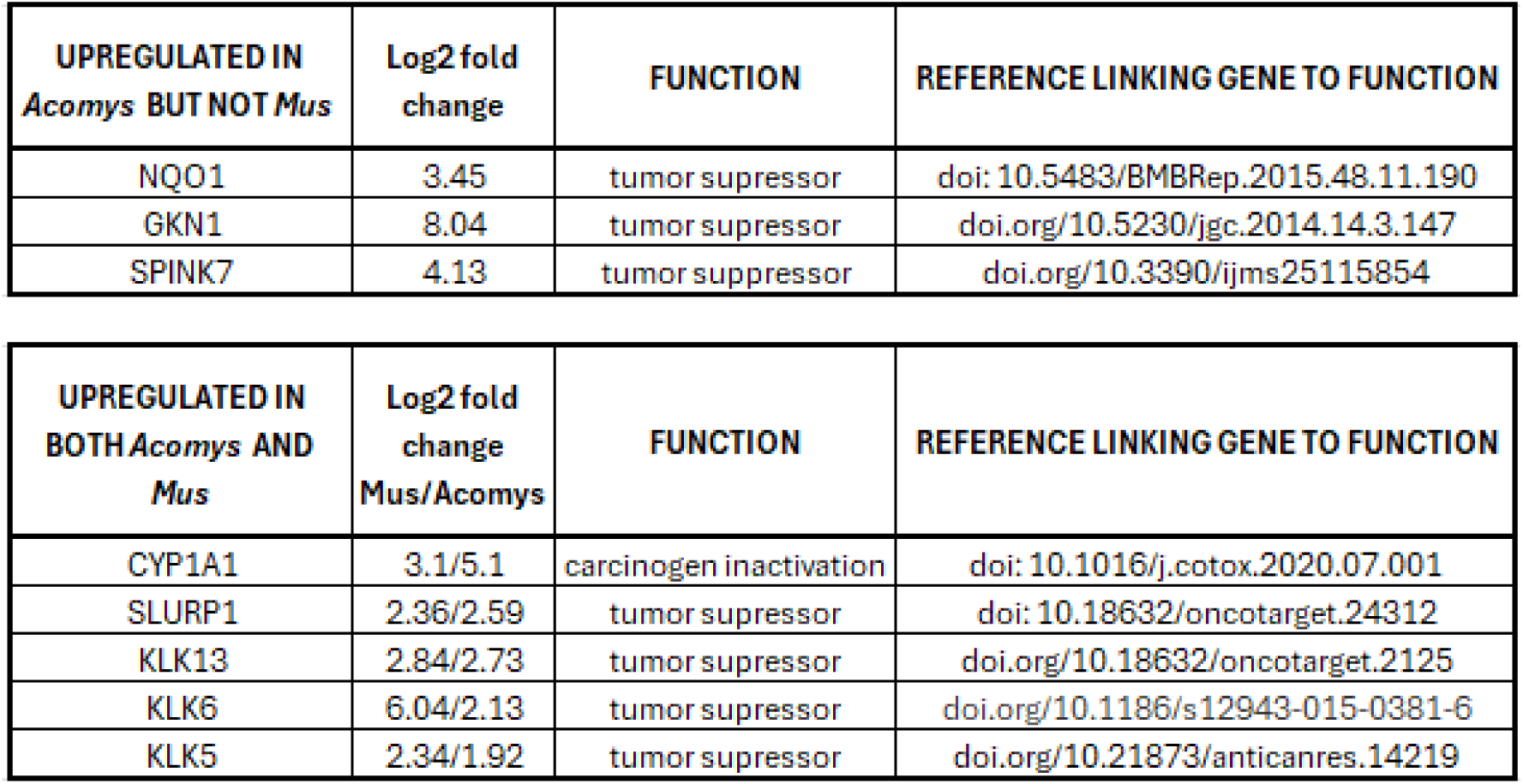
List of genes upregulated in *Acomys* but not *Mus*, or upregulated in both species at D1 after treatment, with references linking them to relevant functions.

### Transcriptional Profile at Day 14

*Mus* and *Acomys* samples at D14 (14 days after treatment with DMBA followed by TPA treatment at D7, D9 and D11) were compared to untreated samples harvested at D0 (N=4). PCA analysis with the expected clustering of samples again showed some heterogeneity for *Acomys* samples, but overall, the PCA1 and PCA2 components explained 53.02% and 19.73% of variation in *Mus* and 41.91% and 23.97% in *Acomys,* respectively (Supplementary Figure 1). We queried DEGs (log2 =2) and p-adj ≤ 0.05 and comparatively examined genes exclusively upregulated in *Mus* but not in *Acomys* (399) and genes exclusively upregulated in *Acomys* but not in *Mus* (303) (Figure 4A, 4B and 4D). Only 39 genes were upregulated in both species (Figure 4B). To understand what BPs were associated with the set of genes upregulated exclusively in each species, we performed a Panther analysis. Genes upregulated exclusively in *Mus* revealed 170 enriched BP ontological categories, while we found 464 BP ontological categories significantly enriched for genes exclusively upregulated in *Acomys*, and only 3 BP ontological categories enriched based on the 33 genes upregulated in both species (Figure 4C). Analysis of the distribution of genes related to the enriched BP categories revealed striking differences in how each species responded to the treatment at this timepoint. In *Mus*, the number of genes in enriched BP categories could be grouped in the following biological themes: cell cycle and proliferation related processes (37.8 %); processes related to epidermis structure and morphogenesis (10.7%); processes related to apoptosis (1.1%) and other processes (50.4%). In contrast, BP categories enriched in

**Figure 4:**
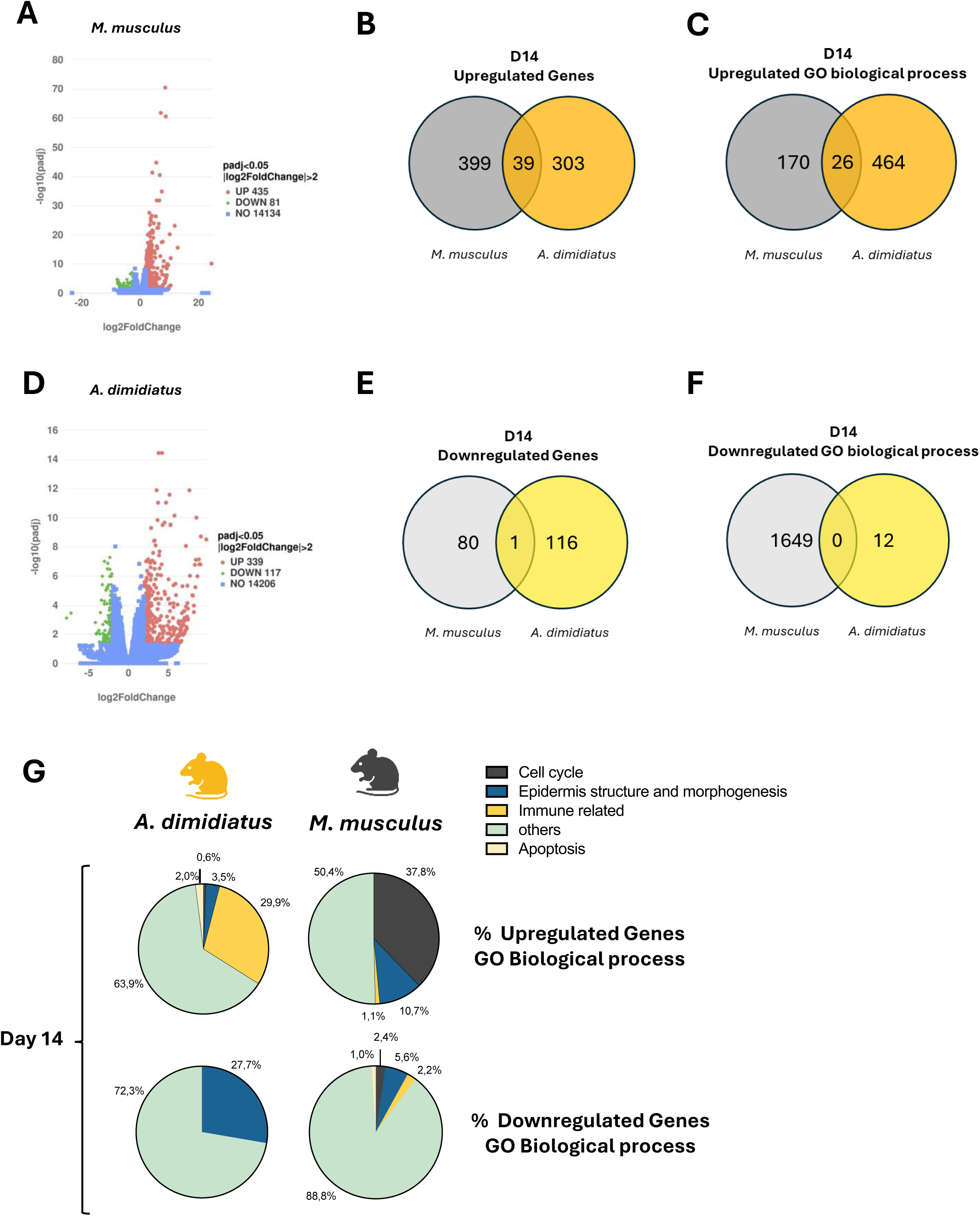
DMBA/TPA treatment induces activation of immune related genes in *A. dimidiatus* versus cell cycle regulation in *M. musculus*. **(A)** and *A. dimidiatus* (D) 14 days after treatment with DMBA (D14). The log2 fold change (FC) indicates the mean expression level for each gene. Genes were scored as differentially expressed when log2 FC>2, p<0,05. Each dot represents one gene. Red dots represent upregulate genes, green dots represent downregulated genes and blue dots represent genes that are not differentially expressed. Venn Diagrams of upregulated genes **(B)** and GO biological processes **(C)**, downregulated genes **(E)** and biological processes **(F)** in both species. **(G)** Percentage of upregulated and downregulated genes included in categories of GO biological processes

*Acomys* were related to immune response (29.9%), apoptosis (2%), cell cycle (0.6%), processes related to epidermic structure and morphogenesis (3.5%) and other processes (63.9%M; Figure 4C and 4G). We examined the identity and known functions of genes upregulated exclusively in *Acomys* vs exclusively in *Mus*. Interestingly, we found that *Acomys* upregulated a total of 29 genes related to tumor suppression, with an average log2 fold increase of 4.31. Among these, several were upregulated to levels higher than log2 fold change = 6: *AHRR* (6.46); *CD89* (7.4); *CLCA2* (6.98); *GKN1* (6.78): *IL25* (8.41); *SHISA* (6.37); *OAS3* (8.84) and *ST18* (6.37). In contrast, we found that *Mus* upregulated only 10 genes related to tumor suppression, and to an average log2 fold level of 3.04 (Table 2).

**Table 2:**
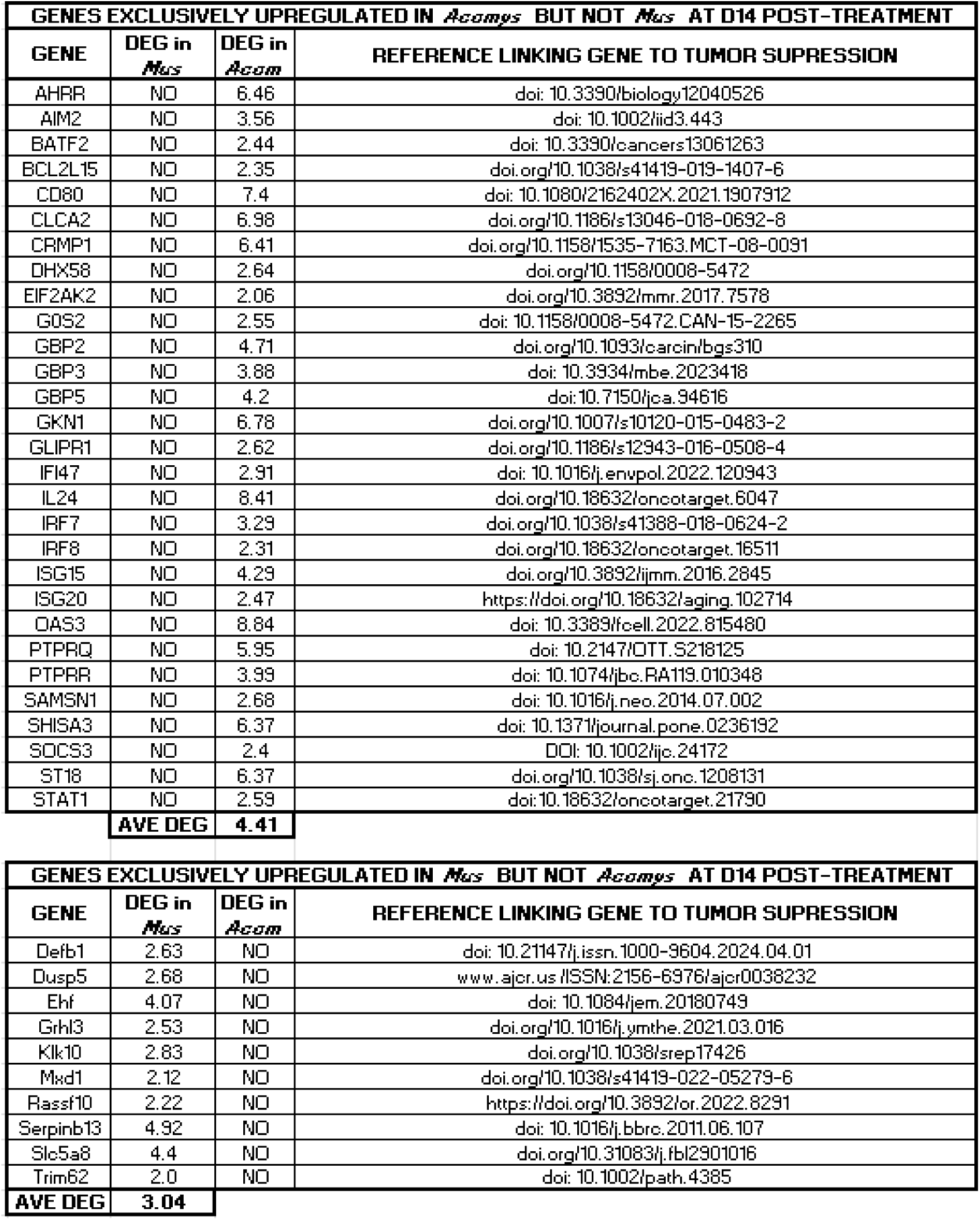
List of genes exclusively upregulated in *Acomys* but not *Mus* related to tumor suppressor functions; List of genes exclusively upregulated in *Mus* but not *Acomys*, related to tumor suppressor functions.

Similarly, we examined genes exclusively downregulated in *Mus* but not in *Acomys* (80) and genes exclusively downregulated in *Acomys* but not in *Mus* (116) (Figure 4A, 4D and 4E). Only 1 gene was downregulated in both species (Figure 4E). When these gene sets were subjected to Panther analysis, a total of 1648 BP ontological categories were found to be enriched in *Mus*, with the associated genes organized into the following themes: processes related to epidermal structure and morphogenesis (5.6%), immune related processes (2.2%), cell cycle and proliferation related processes (5.6%), apoptosis related processes (1%) and other processes (88.8%). Only 12 BP categories were enriched for *Acomys*, 8 of which were related to processes related to epidermal structure and morphogenesis (Figure 4G).

In addition, a BP and KEGG pathway analysis conducted with Novomagic (Novogene Europe) indicated that the response to the DMBA/TPA treatment at D14 did not yield any enriched BP categories or KEGG pathway involving downregulated genes in either *Mus* or *Acomys.* However, the analysis did reveal a number of BP categories and KEGG pathways significantly enriched for upregulated genes in both species (Supplementary Figure 3). Collectively, the data from the D14 analysis suggest that the response in *Mus* primarily involves processes related to the cell cycle and morphological structures while *Acomys* regulates pathways associated with immune response, apoptosis, and tumor suppression.

Given the strong evidence of immune cell infiltration into affected tissues in both species, we conducted IHC against several infiltration markers (Figure 5A). Although, CD45, a pan-leukocyte marker, failed to work in *Acomys* (data not shown), both CD68 and Iba1, which are pan-markers for macrophages revealed different levels of infiltration in *Acomys* vs. *Mus* (Figure 5C, D and E). While these two markers are generally accepted as macrophage markers, it is noteworthy that *Mus* exhibits a higher number of CD68+ cells, while *Acomys* shows higher numbers of Iba1+ cells. This discrepancy suggests that these markers do not identify the same set of cells in both species. Furthermore, IHC against cleaved-caspase 3, an apoptotic marker, showed little or no signal in *Mus*, but showed higher levels in *Acomys* at D14, suggesting an enhanced apoptotic response in *Acomys* at this time point, despite the relatively small number of apoptosis related genes uncovered by our transcriptomic analysis (Figure 5B and E).

**Figure 5.**
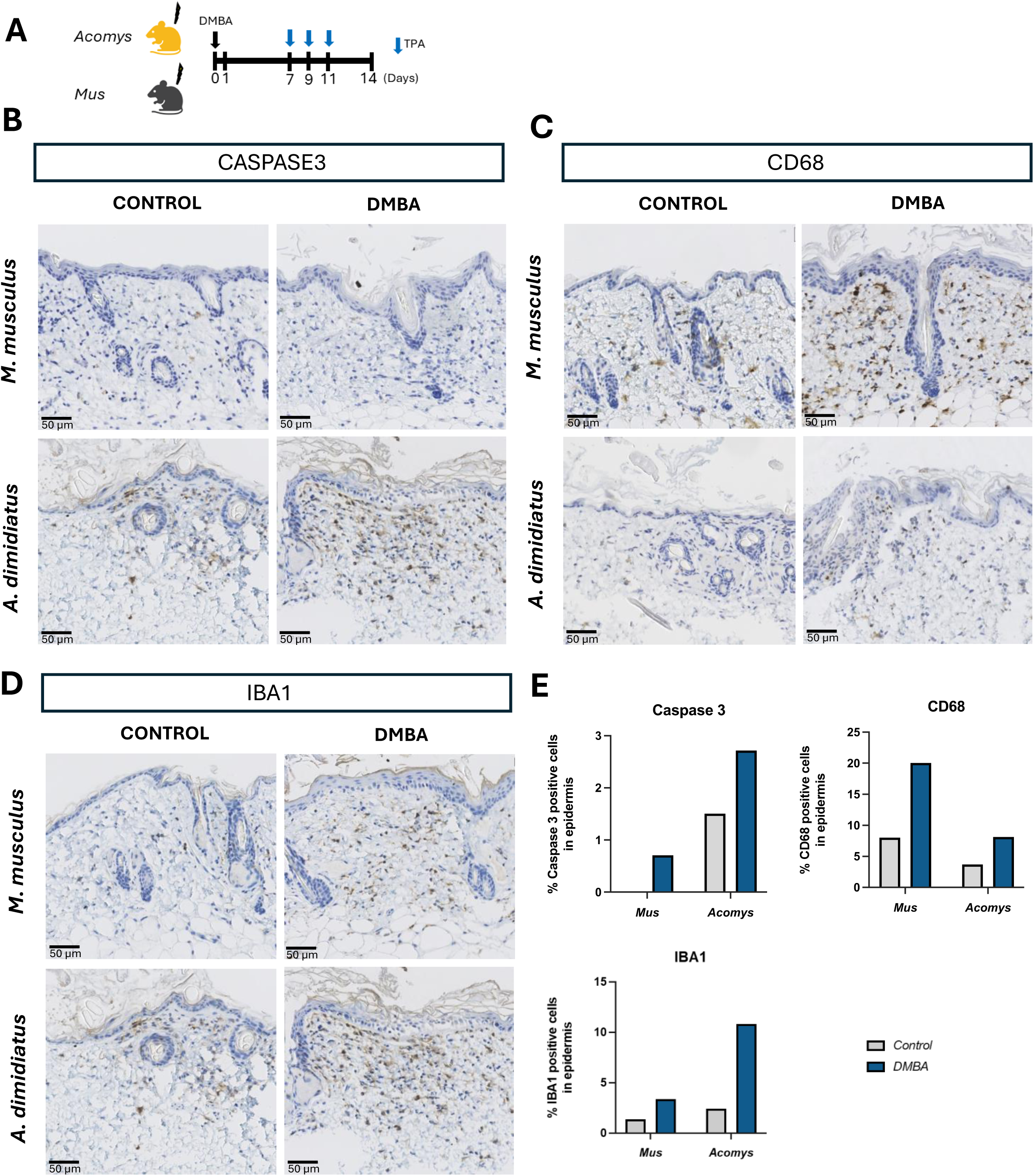
DMBA induces expression of CD68 in *M. musculus* and IBA1 in *A. dimidiatus*, as well as cleaved-Caspase 3 in *A. dimidiatus* at D14. (A) Schematic illustration of the experimental approach to investigate short-term responses to DMBA treatment of both species. **(B-C)** immunohistochemistry staining of **(B)** cleaved- CASPASE3-positive cells, **(C)** CD68-positive cells and **(D)** IBA1-positive cells in *M. musculus* and *A. dimidiatus* at D14 after DMBA/TPA application. **(E)** Quantification of cleaved-caspase 3, CD68 and IBA1 positive cells in both *M. musculus* and *A. dimidiatus*. Scale bar: 50 μm.

### Transcriptional Profile at D28

*Mus* and *Acomys* samples at D28 (after initial treatment with DMBA and TPA treatment three times a week from D7 to D28) were compared to untreated samples harvested at D0 (N=4). PCA analysis showed expected clustering of samples for *Mus*, with PCA1 and PCA2 explaining 77.4% and 11.64% of variability in *Mus*. *Acomys* samples showed one sample clustering with D0 samples, although overall, PCA1 and PCA2 explain 54.76% and 16.96% of the variance (Supplementary Figure 1). We queried differentially expressed genes (log2 =2) and p-adj ≤ 0.05. A total of 1217 genes were exclusively upregulated in *Mus*, while only 46 genes were exclusively upregulated in *Acomys.* Only 3 genes were upregulated in both species (Figure 6A, 6B and 6D). Panther analysis revealed that genes upregulated in *Mus* were associated with 644 significantly enriched BP ontological categories significantly enriched, with these genes relating to BPs in the following proportions: processes related to epidermal structure and morphogenesis (4.9%), processes related to cell cycle and proliferation (10.7%), and other processes (84.2%) (Figure 6C and 6G). No categories were enriched for genes exclusively upregulated in *Acomys* or upregulated genes shared by both species (Figure 6C). Conversely, our analysis of downregulated genes revealed a total of 1763 and 6 genes exclusively downregulated in *Mus* vs *Acomys, respectively* with no genes downregulated in common, and 1528 and 0 processes exclusively downregulated in *Mus* vs *Acomys,* respectively (again with no processes downregulated in common between both species) (Figure 6A, 6D and 6E). When the set of downregulated genes in *Mus* was analyzed using the Panther pipeline, a total of 1528 BP categories were significantly enriched. The genes associated with these BPs corresponded to biological themes in the following proportions: epidermal structure and morphogenesis (2.8%), processed related to cell cycle and proliferation (0.9%), immune related processes (8.5%), apoptosis related processes (1.2%), and other processes (86.5%) (Figure 6F and 6G). In summary, at D28 after initiation of the treatment, *Mus* continues to show significant differential expression of genes, mostly related to cell cycle control and epidermal structure and morphogenesis, but remarkably, the strong differential expression response seen in *Acomys* at D14 (303 and 116 up and downregulated genes respectively) seems to have declined to 57 and 7 up and downregulated genes, respectively, suggesting a return of gene expression almost to baseline levels in *Acomys*. Furthermore, a BP and KEGG pathway analysis conducted with Novomagic (Novogene Europe) shows that response to the DMBA/TPA insult at D28 does not result in either BP categories or KEGG pathway enrichment involving downregulated genes in either *Mus* or *Acomys* but did reveal a number of BP categories and KEGG pathways significantly enriched for upregulated genes in both species (Supplementary Figure 4).

**Figure 6:**
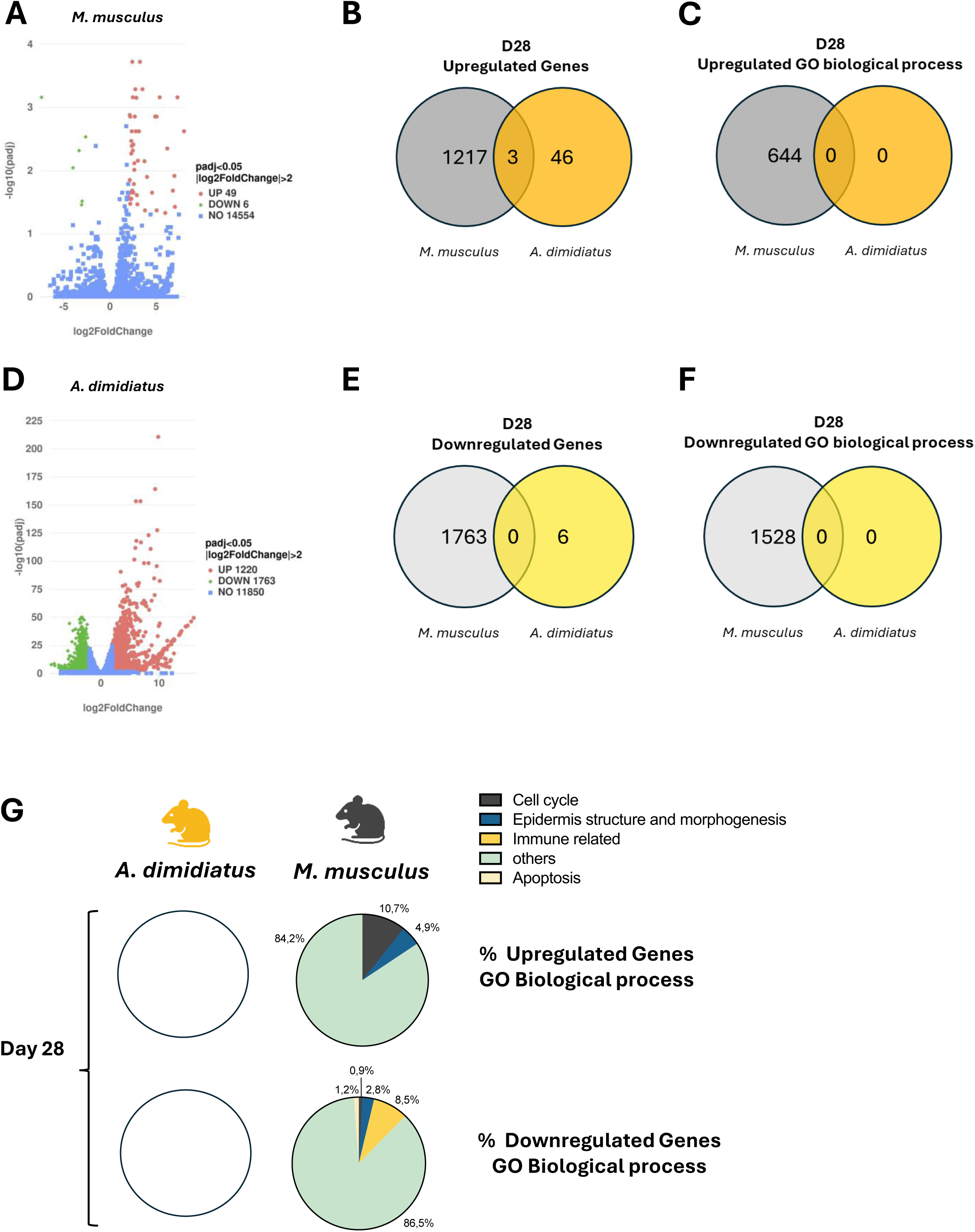
At day 28, *Mus* continues to show significant differential gene expression of genes, but *Acomys* return to baseline. (A, D) Volcano plots of genes differentially expressed in *M. musculus* (A) and *A. dimidiatus* (D) 28 days after treatment with DMBA (D1). The log2 fold change (FC) indicates the mean expression level for each gene. Genes were scored as differentially expressed when log2 FC>2, p<0,05. Each dot represents one gene. Red dots represent upregulate genes, green dots represent downregulated genes, and blue dots represent genes that are not differentially expressed. Venn Diagrams of upregulated genes **(B)** and GO biological processes **(C)**, downregulated genes **(E)** and biological processes **(F)** in both species. **(G)** Percentage of upregulated and downregulated genes included in categories of GO biological processes. White circles are shown when there were no differentially expressed genes identified.

## Discussion

While the African spiny mouse (*Acomys*) is renowned for its remarkable regenerative properties(Maden, Brant, et al., 2018; Maden & Brant, 2018; Matias Santos et al., 2016; Nogueira-Rodrigues et al., 2022; Okamura et al., 2022; Seifert et al., 2012), the relationship between these traits and its incidence and vulnerability to cancer remains largely unexplored. To investigate the sensitivity or resistance to cancer in *Acomys*, we employed a well-established chemically induced carcinogen regimen. This study reports the resistance of *Acomys* to the DMBA/TPA carcinogen-induced tumorigenesis, suggesting enhanced cancer resistance and explores the underlying molecular mechanisms. *Acomys* responds to wounding by mounting a regenerative response, in stark contrast with mammals generally, which repair their injuries by fibrotic scarring (Seifert et al., 2012).;). In response to full thickness 4 mm diameter ear pinna punch wounds (an injury comprising up to 35% of the ear pinna surface), *Acomys* mounts a strong proliferative response causing the tissue (tens of millions of cells) to be re-established and organized into a histological structure that faithfully resembles the original tissue architecture. In contrast, *Mus* simply heals the border of the wound by fibrotic scarring (Matias Santos et al., 2016; Seifert et al., 2012). Dorsal skin wounds in *Acomys* also repair through a regenerative response, re-establishing dermis, epidermis, epidermal appendages, innervation and vascularization, as well as adipose and muscle layers (Sandoval et al., 2020). Acute wound models of *Acomys* of heart and kidney, while falling short of bona fide regeneration, show resistance to ischemic injury (Koopmans et al., 2021; Okamura et al., 2021). Injection of myotoxins into striated muscle, while initially causing severe cellular damage, are undetectable after 2 months (Sandoval et al., 2020). None of these responses are observed in *Mus*. Perhaps most remarkably, a complete spinal cord transection in *Mus* leads to permanent loss of bladder control and hindlimb sensory/motor function. In contrast, *Acomys* with fully transected spinal cords regain bladder control within three weeks and recover up to 60% of motor function within 60 days post-injury (Nogueira-Rodrigues et al., 2022). These observations establish a powerful comparative framework in which two closely related species separated by only 23 million years of evolution show remarkably different responses to wounding (regeneration in *Acomys* vs. fibrotic scaring in *Mus*) in multiple tissues. It is important to recognize that the relationship between enhanced regenerative capabilities and cancer susceptibility is complex, and a deeper mechanistic understanding remains elusive. While regenerative capacities may confer protection against tumorigenesis, rapid cell proliferation, an essential component of regeneration, can also increase the likelihood of mutations and tumor development. Interestingly, a recent report found significant roadblocks to reprogramming *Acomys* cells, suggesting that tumor suppressor pathways play an important role in *Acomys* regeneration and that *Acomys* may possess unreported cancer resistance(Sandoval et al., 2022). Remarkably, over the past decade of research with a *Acomys* colony, which has involved more than a thousand animals, we have consistently observed that *Acomys* initiates a robust proliferative response early in the healing process but never recorded uncontrolled proliferation or the development of spontaneous tumors in aged animals. These observations strongly suggest that regenerative capacity is linked to specific mechanisms that regulate proliferation, potentially leading to cancer resistance as a secondary consequence. This accumulated yet circumstantial evidence prompted us to conduct a systematic evaluation of *Acomys*’ cancer susceptibility. To this end we employed a widely used two-stage carcinogenesis model involving DMBA, a potent carcinogen that induces DNA damage and initiates tumorigenesis and TPA, a tumor promoter that enhances cell proliferation and promotes the progression of initiated cells to cancer. We compared the established response of *Mus* to this insult with that of *Acomys*. Taken together, our data indicate that a) *Acomys* shows resistance to tumor induction using a protocol that clearly induces papillomas in *Mus*; 2) This difference is not due to an initial resistance to DMBA induced DSBs in *Acomys*; 3) Both species mount an initial proliferative response; 4) both species undergo immediate infiltration by immune cells; 4) *Acomys* shows increased apoptotic activity during the first 2 weeks of treatment; 5) the transcriptomic response of *Mus* and *Acomys* to the treatment differ radically and can be summarized as follows: At D1, *Mus* reacts with differential upregulation of gene expression in a total of 414 genes, while *Acomys* upregulates only 17 genes. The BP categories enriched in *Mus* focus mainly on cell cycle and epidermal structure and morphogenesis. In contrast, the 17 genes upregulated in *Acomys* include upregulation of the detoxification gene *NQ01*, as well as other genes with cancer prevention functions, including the tumor suppressor *SPINK 7*; at D14, the transcriptomic response is significant in both species in terms of number of DEGs, but radically distinct in terms of the functions those genes are involved in.

Our data clearly show that by D14, the response of *Mus* to the genotoxic insult focuses on modifications to epidermal structure and an attempt to modulate proliferation and cell cycle related processes, while *Acomys*, in contrast, unleashes a powerful and multifaceted immune response. This suggests that the molecular strategies against tumorigenesis are radically different in these two species, and that *Acomys sp* may possess unique mechanisms to resist tumorigenesis, even under conditions that typically induce cancer in non-regenerative mammals. We establish, for the first time, a tractable and logistically convenient comparative model to specifically explore cancer resistance mechanisms in vivo in the laboratory which may have significant implications for understanding the links between regeneration and cancer resistance and offer novel insights into potential therapeutic approaches to limit tumor development in humans.

These results fit the pattern of animal model studies found in the literature associating high regenerative capabilities to cancer resistance, in this instance in a rodent phylogenetically close to humans. It is tempting to speculate that the key relation between regeneration and cancer resistance resides in the ability to control and cease proliferation during wound healing in an adaptive manner, and that these mechanisms are supercharged in a species that uses regeneration as a wound healing strategy. However, our transcriptomic analysis suggests that there are several layers to mechanisms deployed to surveil proliferation in *Acomys,* with the immune system playing a key role in controlling tumorigenic processes. This is evident from the observation that *Mus* seems to concentrate its control efforts on modulating the structure and state of the epidermis (keratinocyte proliferation and differentiation) and controlling cell cycle progression, while *Acomys* modulates 29.1% of the genes it upregulates at D14 of treatment in BP categories related to immune response (Calve et al., 2010; Godwin et al., 2006; Imokawa et al., 2003; King et al., 2012; Levi et al., 2008; McGann et al., 2001; Mescher et al., 2005; Pajcini et al., 2010; Seifert et al., 2012; Tanaka et al., 1999). In the context of the relationship between regeneration and cancer resistance, our results prompt us to focus on the role of the *Acomys* immune system. While there is general consensus in the literature regarding the significance of the *Acomys* immune system in regeneration and several studies have explored specific aspects(Gawriluk et al., 2020; Simkin et al., 2017), much remains unknown. It has been speculated that regeneration in *Acomys* correlates with a blunted immune response involving a decrease in inflamatory cytokines. Gawrilik et al. observed that regeneration was associated with lower levels of pro-inflammatory cytokines (i.e., IL-6, CCL2, and CXCL1) and an increase in local levels of IL-12 and IL-17, correlating with an influx of T-cells into the wound area (Gawriluk et al., 2020). At day 14, we did not find differential expression of IL-6, CCL2, and CXCL1, but identified lower levels of other cytokines considered generally pro-inflammatory, such as IL-12, Il-17, IL-1a and IL-18 in *Acomys*. Also, macrophages have been shown to be required for epimorphic regeneration (Simkin et al., 2017). Our Panther analysis of BP categories enriched in *Acomys* at D14 found that over 50% of BP categories related to a range of processes related to immunity, including T-cell proliferation, differentiation, migration, and cytokine production, NK chemotaxis and differentiation, macrophage associated processes and a number of cytokine-associated pathways. Our results suggest the immune system of *Acomys* is crucially positioned to determine the outcomes of wound healing, regeneration and cancer resistance. The general characterizations of immune responses in *Acomys* as simply blunt or heightened in comparison to that of non-regenerators seem to be premature and fail to capture the complexity and context-dependent response of the system. A thorough dissection of the *Acomys* immune response in relation to regeneration and cancer resistance is likely to keep researchers busy for a long time to come.

## Acknowledgements

This work was supported by a grant from Spanish Ministerio de Ciencia e Innovación/Agencia Estatal de Investigación and the European Regional Development Fund (PID2022-136654OB-I00 financed by MCIN/AEI /10.13039/501100011033 / FEDER, UE). This study received Portuguese national funds from FCT - Foundation for Science and Technology through projects UIDB/04326/2020 (DOI:10.54499/UIDB/04326/2020), UIDP/04326/2020 (DOI:10.54499/UIDP/04326/2020) and LA/P/0101/2020 (DOI:10.54499/LA/P/0101/2020). We acknowledge the expertise and dedication of the Animal House of ABC Ri-UAlg.

## Supplementary Figures Legends

**Supplementary** Figure 1 **- Principal component analysis (PCA) for *Acomys dimidiatus* and *Mus musculus* samples**. **(A)** PCA analysis of samples collected one day after application of DMBA in both *A. dimidiatus* and *M. musculus*. **(B)** PCA analysis of samples collected 14 days after application of short-term DMBA/TPA protocol in both *A. dimidiatus* and *M. musculus*. **(C)** PCA analysis of samples collected 28 days after application of short-term DMBA/TPA protocol in both *A. dimidiatus* and *M. musculus*. MM_D0 represent M. musculus samples at day 0, AC_D0 represent *A. dimidiatus* samples at day 0, MM_D1_DMBA represent *M. musculus* samples collected 1 day after treatment with DMBA, AC_D1_DMBA represent *A. dimidiatus* samples collected 1 day after treatment with DMBA, MM_D14_DMBA represent *M. musculus* samples collected 14 days after treatment with DMBA/TPA, AC_D14_DMBA represent *A. dimidiatus* samples collected 14 days after treatment with DMBA/TPA, MM_D28_DMBA represent *M. musculus* samples collected 28 days after treatment with DMBA/TPA, AC_D28_DMBA represent *A. dimidiatus* samples collected 28 days after treatment with DMBA/TPA.

**Supplementary** Figure 2 **- DMBA treatment upregulated several GO biological processes and Kegg pathway. (A)** GO Biological processes upregulated and downregulated in *Mus musculus* and *Acomys dimidiatus*, 24h after the treatment with DMBA. **(B)** Kegg pathways upregulated and downregulated in *M. musculus* and *A dimidiatus*, 24h after the treatment with DMBA.

**Supplementary** Figure 3 **- DMBA treatment upregulated several GO biological processes and Kegg pathway. (A)** GO Biological processes upregulated and downregulated in *Mus musculus* and *Acomys dimidiatus*, 14 days after the treatment with DMBA. **(B)** Kegg pathways upregulated and downregulated in *M. musculus* and *A dimidiatus*, 14 days after the treatment with DMBA.

**Supplementary** Figure 4 **- DMBA treatment upregulated several GO biological processes and Kegg pathway. (A)** GO Biological processes upregulated and downregulated in *Mus musculus* and *Acomys dimidiatus*, 28 days after the treatment with DMBA. **(B)** Kegg pathways upregulated and downregulated in *M. musculus* and *A dimidiatus*, 28 days after the treatment with DMBA.

