## Supplementary figures and images for "Resistance to Carcinogenesis in the African Spiny Mouse (*Acomys*) correlates with upregulation of tumor suppressor genes"

### Supplementary Figures

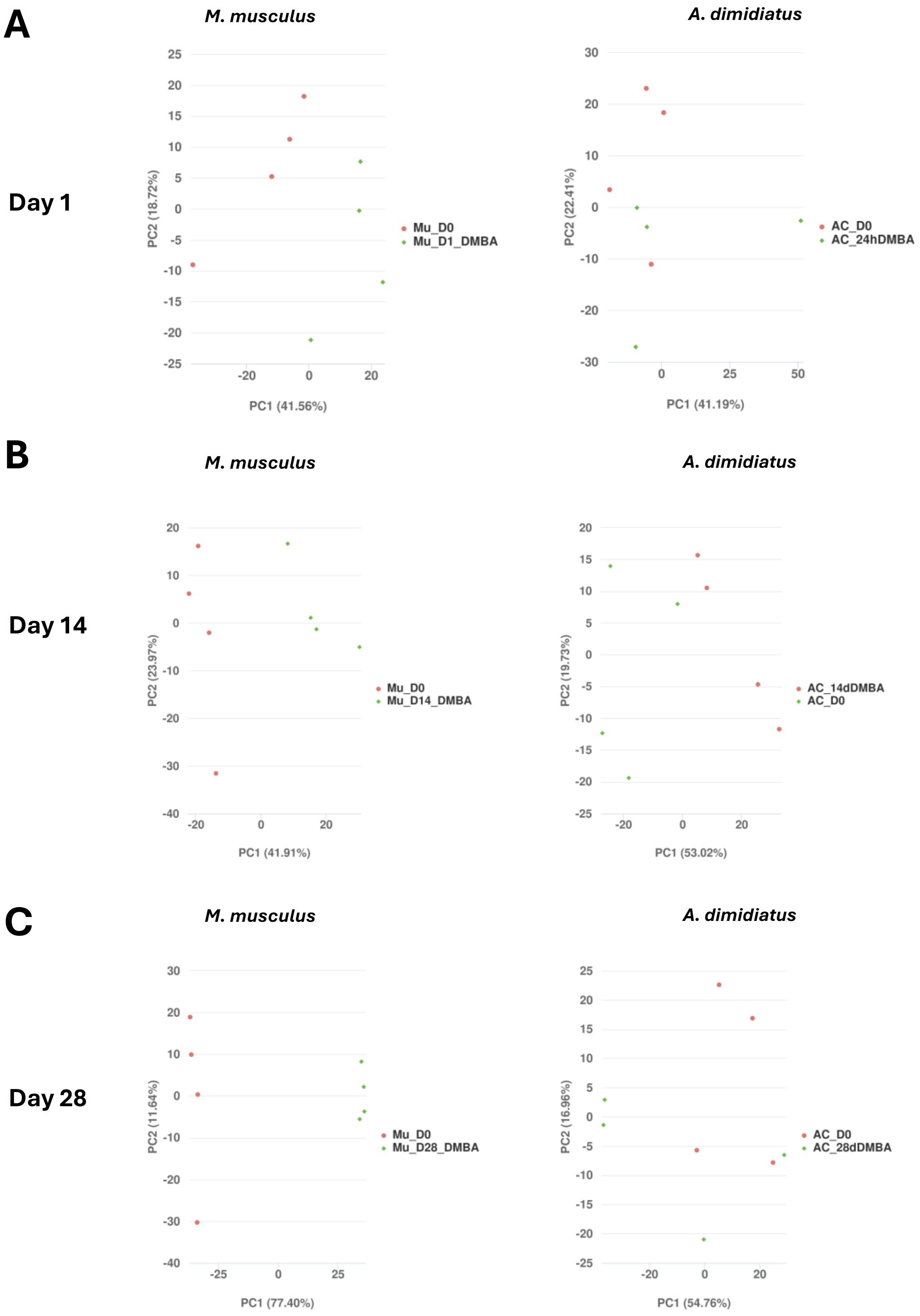

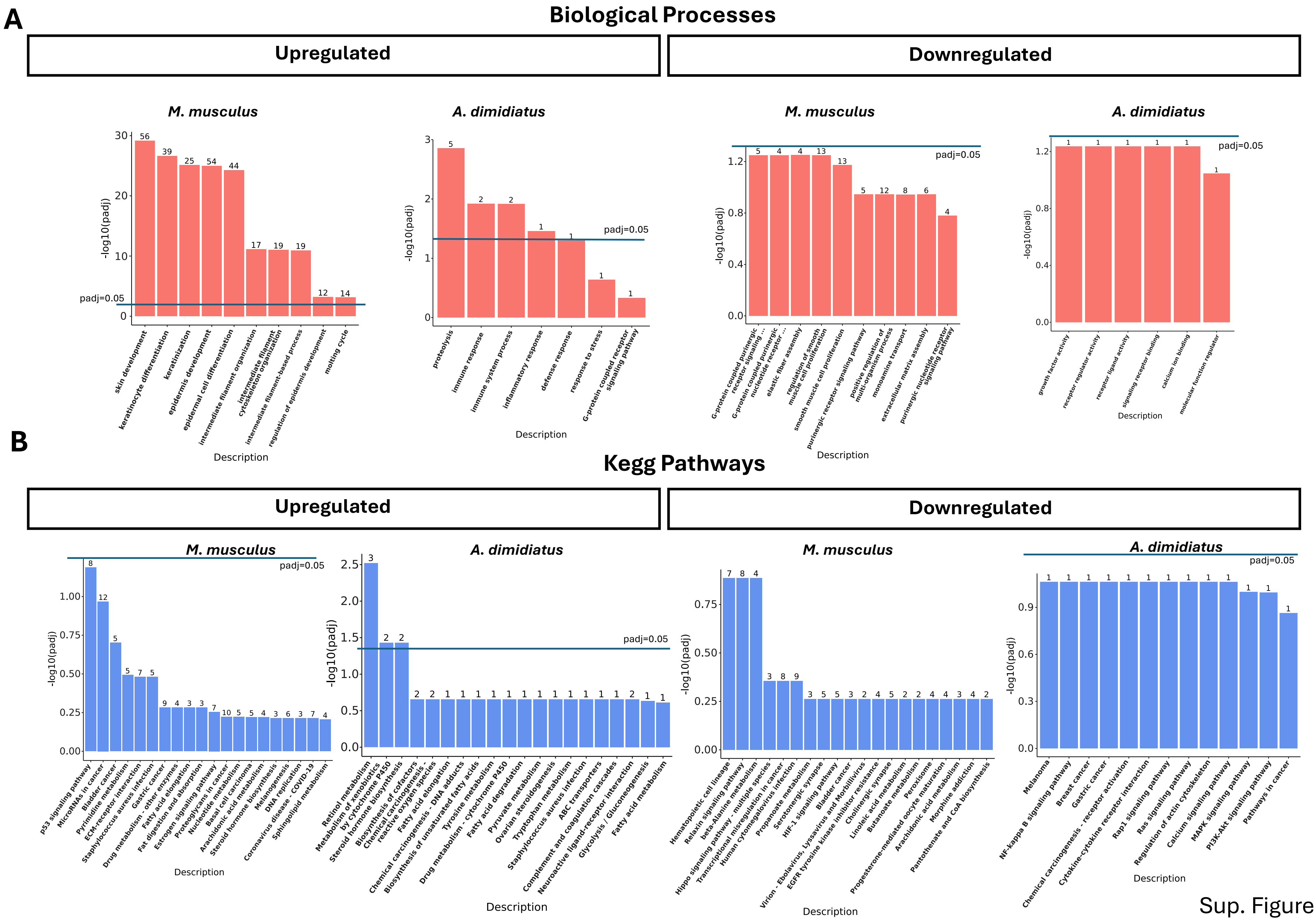

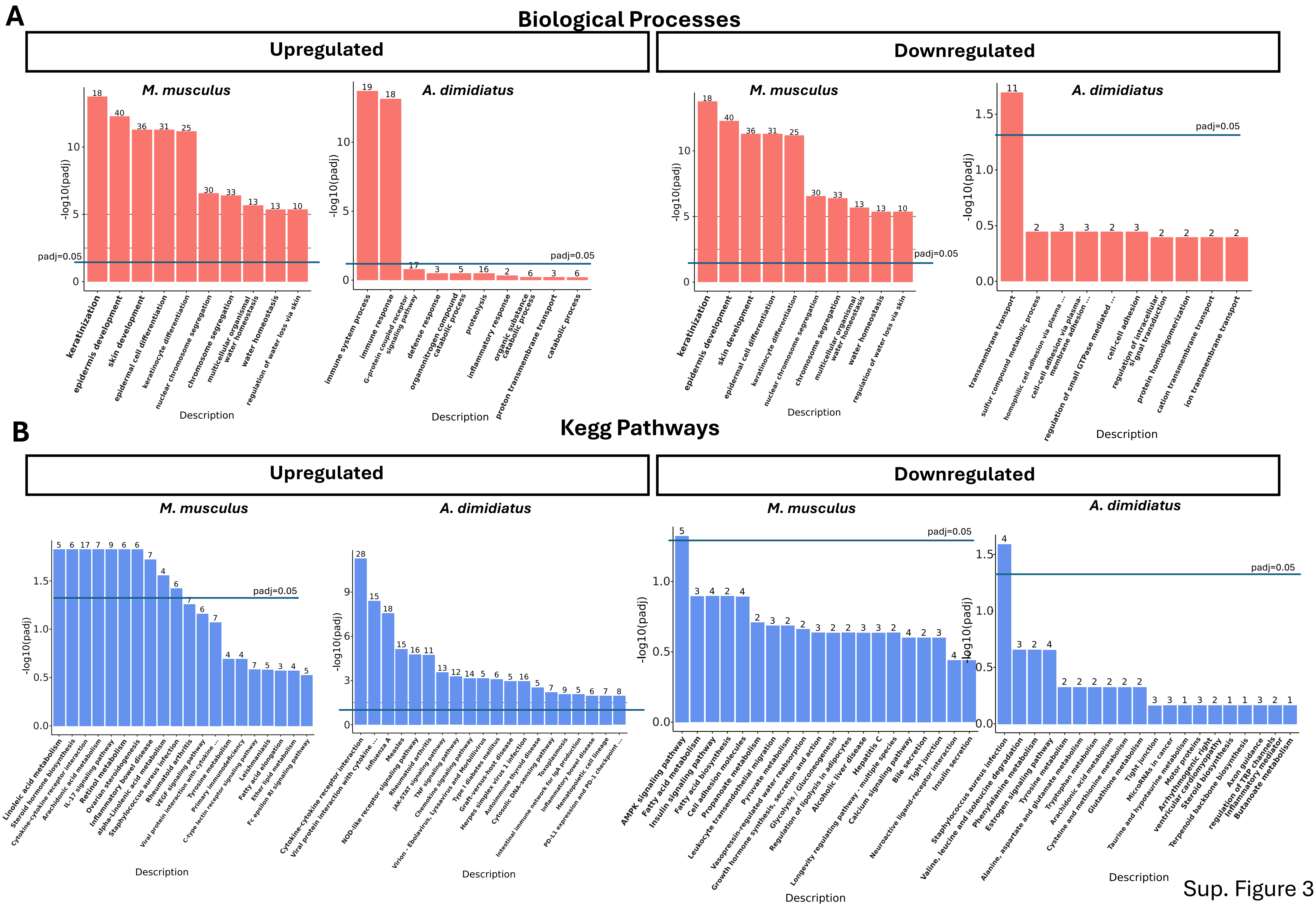

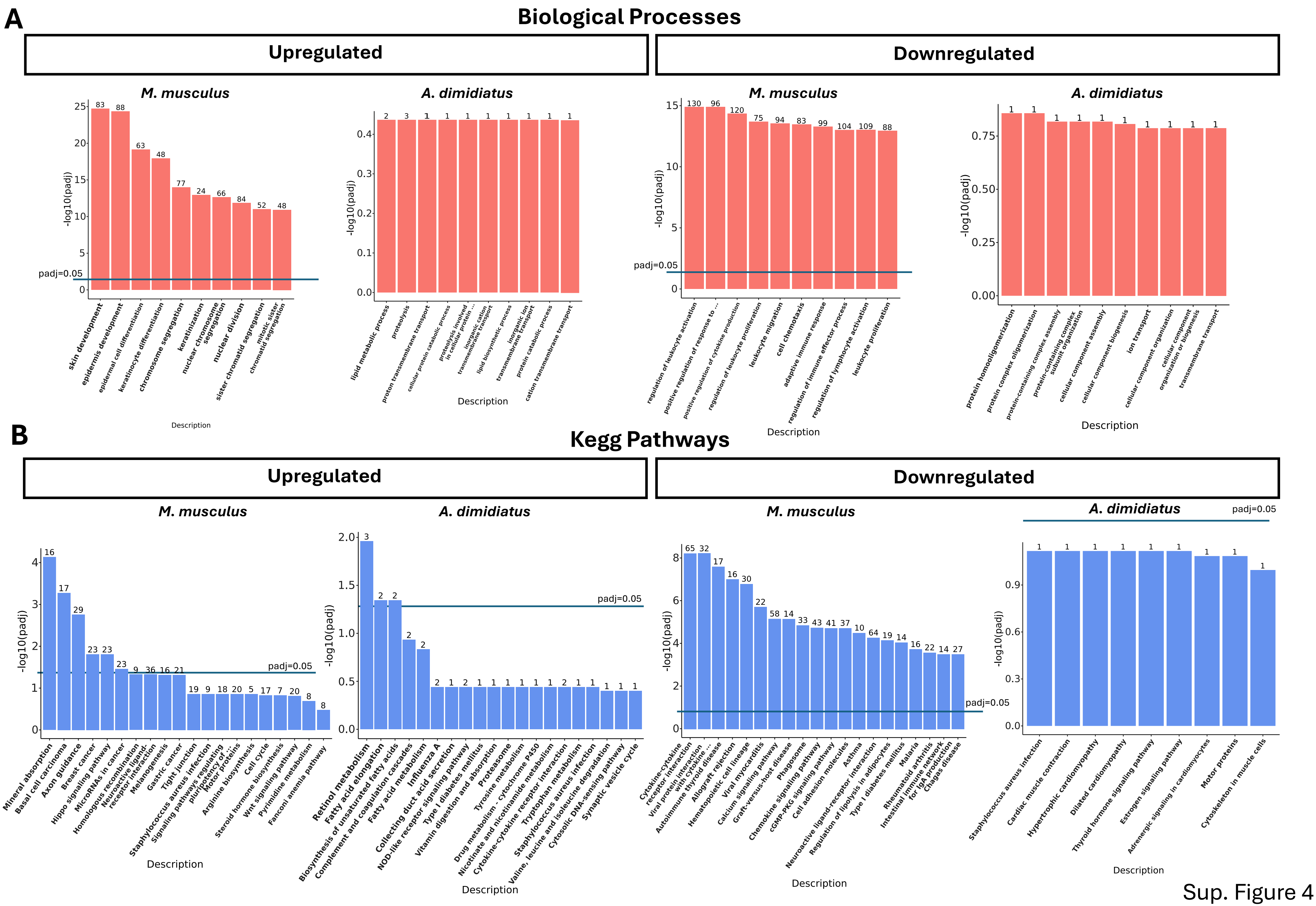
